# Egg-laying hormone expression in identified neurons across developmental stages and reproductive states of the nudibranch *Berghia stephanieae*

**DOI:** 10.1101/2023.12.21.572887

**Authors:** Cheyenne C. Tait, M. Desmond Ramirez, Paul S. Katz

## Abstract

Neuropeptides play essential roles in coordinating reproduction. Egg-laying hormone (ELH) is conserved in genetic sequence and behavioral function across molluscs, where neuronal clusters secrete ELH to modulate and induce egg-laying. Here we investigated ELH in the nudibranch mollusc, *Berghia stephanieae*. ELH preprohormone gene orthologs, which showed clade-specific differences at the C-terminus of the predicted bioactive peptide, were identified in brain transcriptomes across several nudipleuran species, including *B. stephanieae*. Injection of synthesized *B. stephanieae* ELH peptide into mature individuals induced egg-laying. ELH gene expression in the brain and body was mapped using *in-situ* hybridization chain reaction. Across the adult brain, 300-400 neurons expressed ELH. Twenty-one different cell types were identified in adults, three of which were located unilaterally on the right side, which corresponds to the location of the reproductive organs. Ten cell types were present in pre-reproductive juvenile stages. An asymmetric cluster of approximately 100 small neurons appeared in the right pedal ganglion of late-stage juveniles. Additional neurons in the pleural and pedal ganglia expressed ELH only in adults that were actively laying eggs and sub-adults that were on the verge of doing so, implicating their direct role in reproduction. Outside the brain, ELH was expressed on sensory appendages, including in presumptive sensory neurons. ELH shares deep homology with the corticotropin-releasing hormone gene family, which has roles broadly in stress response. Its widespread expression in the nudibranch *B. stephanieae* suggests that ELH plays a role beyond reproduction in gastropod molluscs.

**Highlights:**

- Egg-laying hormone (ELH) preprohormone sequences were identified in the transcriptomes of several nudipleuran molluscs.
- Many ELH-expressing neurons and clusters in the brain could be individually identified based on soma position and morphology.
- Some identified ELH-expressing neurons were not observed until later juvenile stages.
- Some neurons expressed ELH in adults only when they were actively laying eggs.
- Many hundreds of ELH-expressing neurons were present in peripheral appendages.

## Introduction

Neuropeptides are ancient signaling molecules showing deep homology across the tree of life whose functions are often multilayered and pleiotropic (Odekunle and Elphick, 2020; Thiel et al., 2021). In gastropods, one of the most well-studied neuropeptides is egg-laying hormone (ELH), possessing sequence conservation and conservation of bioactivity (Hermann et al., 1997; Mahon and Scheller, 1983). Release of the aptly named neuropeptide from well-characterized secretory neurons results in a stereotyped sequence of egg-laying behaviors. Here, we identified the ELH sequence in the nudibranch *Berghia stephanieae*, verified its activity and localized its expression in the brain and peripheral tissues across developmental and reproductive states. In the future, we can study its roles throughout the brain and body.

Egg-laying hormone was initially discovered in the sea hare *Aplysia californica* (Arch, 1976). This was followed by the discovery of its ortholog, the caudo-dorsal cell hormone (CDCH), in the pond snail *Lymnaea stagnalis*, which similarly induced egg-laying behaviors (Roubos, 1984). Similarities were seen in cellular morphology and electrophysiological characteristics of the secretory neurons producing these hormones in each species, with these cell clusters termed bag cells in *A. californica* and caudo-dorsal cells (CDCs) in *L. stagnalis*. Sexual maturity is synchronized with the appearance of the bag cells and CDCs, underscoring the involvement of these neurons in reproduction (De Lange et al., 1994; McAllister et al., 1983).

However, there are indications that ELH expression may not be limited to clusters of secretory neurons inducing egg-laying behavior in mature animals. *A. californica* has over 20 ELH-expressing neurons in all regions of the brain except the pedal ganglia, consistently found across developmental stages (McAllister et al., 1983). *L. stagnalis* also has dozens of ELH-expressing neurons in stereotyped locations, with some in the pleural ganglia appearing less stereotyped in their position (van Minnen et al., 1988). In *A. californica*, some clusters in the central ganglia are similar in their electrophysiological characteristics to the bag cells, and may be coupled with them, forming a circuit of descending control of egg-laying behaviors (Brown et al., 1989; Hatcher and Sweedler, 2008). There are additional neurons expressing ELH that are not part of these clusters, and whose actions and identities have not been documented in these systems.

Although conserved across molluscs including gastropods and bivalves (De Oliveira et al., 2019), there has been no indication of ELH in the nudipleuran gastropods despite recent multi-species transcriptomics studies (Lee et al., 2021). However, there is evidence for the existence of an ELH-like hormone. In *Pleurobranchaea californica* injection of homogenized cerebropleural ganglia from an actively laying individual induces egg-laying (Ram et al., 1977). Also, antisera for *A. californica* ELH labeled some neurons in the cerebral ganglion of the nudibranch *Archidoris montereyensis* (Wiens and Brownell, 1994). Here, we used brain transcriptomes to identify ELH gene orthologs in several nudipleuran species. We characterized the neuronal gene expression of ELH in the brain and periphery of one member of this group, the nudibranch *Berghia stephanieae,* and determined its developmental and state-dependence.

## Material and Methods

### Animals

*Berghia stephanieae* was obtained commercially from Salty Underground (Missouri, USA) and reared communally in groups of up to 30 animals, in artificial sea water (ASW, Instant Ocean) on a 12:12 LD cycle with aeration provided by an air stone. Every other day, one live, large *Exaiptasia pallida* anemone was placed in each tank as food. As reported by other research groups, in this environment individuals mated multiply and laid up to 4-6 egg masses per week (Monteiro et al., 2020). The feeding and light regimes were continued during all long-term behavioral observations.

### Brain transcriptome generation and assembly

Generation of transcriptomes utilized previously published RNA sequencing data for *Melibe leonina* (Cook et al., 2018), *Hermissenda crassicornis* (Tamvacakis et al., 2015), *Pleurobranchaea californica* (Tamvacakis et al., 2018), *Dendronotus iris* (Ramirez et al., 2023), and *Berghia stephanieae* (Ramirez et al., 2023). For *Flabellinopsis iodinea*, 26 animals (0.9-2.4 grams) were obtained from Living Elements (Delta, Canada), Monterey Abalone Company (Monterey CA), and Marinus Scientific LLC (Newport Beach, CA). Animals were kept in recirculating artificial seawater aquaria at 10-12°C on a 12:12 light/dark cycle. *Armina spp.* was similarly purchased and maintained prior to dissection. Dissected brains for each species were flash-frozen and pooled into respective 1.5 mL centrifuge tubes, and total RNA was extracted using a QIAGEN RNeasy Plus Universal Midi Kit. RNA quality was assessed with a Bioanalyzer 2100 (Agilent) and sent to Beckman Genomics where messenger RNAs were isolated from each sample using poly-A capture, converted to cDNA, and Illumina sequencing adaptors were ligated for sequencing on an Illumina HiSeq 2500.

After sequencing, and for previously published data, we used the Oyster River Pipeline (ORP) to assemble reads into a transcriptome for each species, as previously completed for *B. stephanieae* and *D. iris* (Ramirez et al., 2023). Briefly, the Oyster River Pipeline (MacManes, 2018) created multiple transcriptomes that were merged and then filtered using OrthoFinder2 (Emms and Kelly, 2019).

### Molecular identification of egg-laying hormone across the nudipleuran sea slugs

We used HMMER (Eddy, 2011) to build a profile from known molluscan ELH peptide sequences, which included both bivalves and gastropods (Supplemental Table 1). We then used HMMER and this profile to search for peptides orthologous to the ELH peptide across our nudipleuran brain transcriptomes in translated form.

In parallel, we used Blastp (Johnson et al., 2008) and the bioactive ELH sequence of *Aplysia californica* to search the same translated nudipleuran transcriptomes. Relaxed e-value thresholds (1.0) and word size (2) were used in consideration of the short (<50 AA) length of these neuropeptides. ELH orthologs from some species *(Melibe, Dendronotus*) were recovered using Blastp, but most were not. To verify our HMMER results, we then used Blastp in an iterative fashion, with the *Dendronotus* putative ELH sequence, and then the *B. stephanieae* putative ELH sequence as our queries (Supplemental Table 2).

Preprohormone amino acid sequences were annotated as in previous studies of ELH in other molluscs (Stewart et al., 2014). We noted the presence of signal peptides using SignalP 6.0 with the molluscan and “long” analysis options selected (Teufel et al., 2022). We also annotated the high probability proteolytic cleavage sites using the eurkaryotic settings of NeuroPred (Southey et al., 2006), reporting the full amino acid sequences of each putative ELH preprohormone with these characteristics annotated (Supplemental Table 3). The bioactive egg-laying hormone peptide was also aligned across species using MUSCLE (Edgar, 2004), then annotated using Jalview (Waterhouse et al., 2009). A tree for the short bioactive ELH amino acid sequence was generated using maximum likelihood methods and bootstrapping with IQTree’s default parameters (Minh et al., 2020), and the final tree generated using ITOL (Letunic and Bork, 2021).

### Observation of spontaneous egg-laying behavior

It is critical to use animals healthy enough and in the right state to lay eggs for these experiments (Stewart et al., 2016). To be certain of egg-laying status, we separated *B. stephanieae* into jars individually and ensured that all individuals laid at least one egg mass within 7 days. To observe spontaneous egg-laying behavior, we transferred 12-20 of these animals into four groups, each within its own one-gallon observation tank, and video recorded them continuously for 5 days. One group of 12 was recorded for only 3 days, for a total of 44 animals observed for 120 hours, and 12 animals observed for 72 hours. Feeding was essential – pilot experiments showed that animals that were not fed did not lay eggs. Thus, all groups were fed 1-2 anemones every day at the same time. The tank was videoed from above with a webcam (Logitech 615) that was modified to be capable of picking up infrared (IR) illumination. The tank was placed on top of a lightboard so that it was backlit during daylight hours. Overnight, it was instead lit from the side by an infrared LED floodlight.

To capture this behavior in greater detail, other groups of animals were watched periodically throughout the day. As soon as an animal was noted to have begun to show counterclockwise turning, they were moved to a Leica M165C dissection microscope equipped with a 10 megapixel CMOS camera (Leica IC90E), where video could be captured using a 0.8X objective, and with a 30 frames per second temporal resolution.

### *B. stephanieae* ELH injection assay

The nucleotide sequence we identified in *B. stephanieae*’s transcriptome for the bioactive ELH peptide was translated into a predicted protein and synthesized by Genscript (Piscataway, NJ, USA) to 95 % purity, with C-terminal amidation. It was diluted in distilled water to 10 ^-4^ M, then to 10^-5^ M in ASW.

For ELH peptide or control injections, animals were isolated in 8 ounce, BPA-free, food-grade, PET plastic jars 3-7 days prior to the injection, to ensure they were capable of laying eggs. All animals used in subsequent injections were observed to have laid at least one egg mass at least 48 hours prior to ELH injections. They had also been fed within the prior two days. Animals were injected into their foot with 3-5 µL of 10^-5^ M ELH using a 10 µL Nanofil syringe (World Precision Instruments) equipped with a 35-gauge needle. This concentration was chosen based on a study of synthesized ELH being injected into another gastropod, *Theba pisana* (Stewart et al., 2016). Control animals were injected with 3-5 µL of ASW using the same procedure. Animals were then videoed in 6-well culture dishes on a lightboard with an overhead webcam, so that they were backlit, for 6-8 hours. They were then moved individually to plastic jars for 24 hours so that any further egg masses could be attributed to injected control or injected ELH animals.

### ELH *in-situ* hybridization chain reaction

Before we could localize expression of the ELH preprohormone, the tissue had to be prepped. First, brains or peripheral appendages were dissected from adults that had been anaesthetized in 4.5% MgCl_2_ in artificial seawater (ASW) on ice for 15 minutes. This tissue and whole bodies of anaesthetized juveniles were fixed at room temperature for 2-3 hours in 4% paraformaldehyde (PFA) in ASW. For the state dependent aspect of this study, adults were observed periodically throughout the day such that they could be captured in the act of spontaneous egg-laying behavior. Individuals found to have laid at least two loops of eggs in a whorl were then interrupted, anaesthetized for 10 minutes in ice cold ASW, then dissected and fixed, with brains dissected free from their tissue before commencing the labeling protocol. Animals labeled during mating were similarly interrupted 1-2 hours into the behavior and fixed in PFA, with brains dissected free of other tissues only after fixation.

To visualize expression of ELH in specific neurons in the brain and in peripheral tissues of *B. stephanieae*, we used in-situ hybridization chain reaction (HCR) (Choi et al., 2018). With the complete mRNA sequence for the *B. stephanieae* egg-laying hormone precursor (reported in the results below and Supplemental Table 4), a library of 30 HCR probe pairs was generated by Molecular Instruments (Los Angeles, California) and maintained as a stock solution at 1 µM.

Briefly, our labeling protocol is as follows: Fixed tissues were incubated in pre-hybridization buffer (Choi et al., 2018) for 30 minutes at 37 °C, with urea substituted for formamide (Sinigaglia et al., 2018). Then they were incubated in 1 µL / 100 µL of ELH probe in hybridization buffer, for a final concentration of 0.01 M probe, for 24 hours at 37 °C. After washing in wash buffer, again with urea substituted for formamide, hairpins complementary for the probes with fluorophores at the 647 wavelength (Molecular Instruments) were added for 24 hours. Hairpins were then washed off with 5x SSCT, brains were cleared with deepclear (Pende et al., 2020) and mounted on glass slides in Vectashield anti-fade medium (Vector Laboratories). Peripheral tissues and juveniles were instead cleared using a fructose-glycerol solution (Dekkers et al., 2019), and then mounted in the same.

### Imaging and image quantification

Images of HCR with the ELH probes were taken at 10x or 20x using an air objective on either a Zeiss 710M LSM or Nikon A1R25 laser-scanning confocal microscope at the Light Microscopy Facility at University of Massachusetts Amherst. ZenBlack (Zeiss) or NIS-Elements (Nikon) software was used to capture the images. These raw z-stacks were then processed, and z-projections made, using FIJI (Schindelin et al., 2012).

## Results

### Nudipleuran ELH amino acid sequences have diverged from other molluscs

We used a bioinformatics approach to identify the ELH preprohormone sequence in *B. stephanieae* and six other nudipleuran species. A protein model was generated in HMMER using only the reported 37-41 amino acid sequences of the bioactive ELH peptide in other molluscs (Supplemental Table 1). This model was then used to scan for and find orthologous sequences in all nudipleuran transcriptomes examined. These sequences were verified using an iterative Blastp approach (see Methods and Supplemental Table 2).

The preprohormone lengths for the nudipleurans ranged from 374 amino acids in *B. stephanieae* to 190 amino acids in *Melibe leonina* (Figure 1A, Supplemental Table 3). For each putative ELH preprohormone ortholog, we identified the signal peptide and potential proteolytic cleavage sites. Three of the seven sequences lacked a signal peptide at the N-terminus, indicating that they were partial sequences. For all seven, the bioactive ELH peptide was present at the C-terminus, which is a conserved structural motif across gastropod molluscs. Each had a predicted proteolytic cleavage site immediately before this bioactive region. There was also a consensus glycine at the C-terminal end of the peptide, which leads to C-terminal amidation when the protein is processed. The total number of predicted proteolytic cleavage sites was highly variable across the nudipleuran species ranging from three in *Hermissenda crassicornis* to eleven in *B. stephanieae*. Such variation is shared in other gastropod clades.

**Figure 1:**
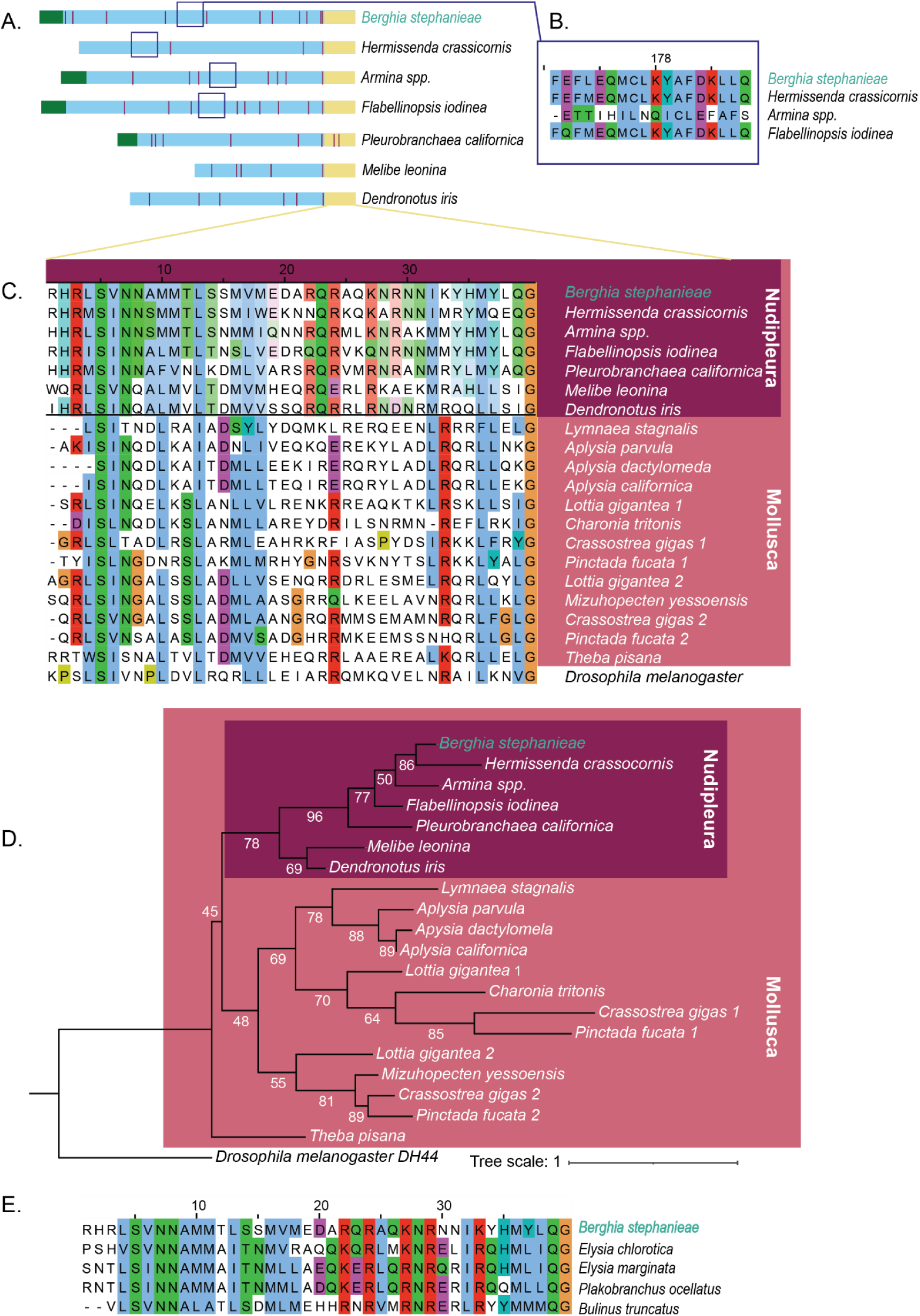
Molecular identification of ELH in the nudipleuran sea slugs. **A.** Preprohormone models for the newly reported nudipleuran sequences show that the overall structure of the ELH gene, with a signal peptide at the N-terminus (green) and the bioactive 37-40 amino acid ELH peptide at the C-terminus (yellow), and proteolytic cleavage sites throughout (purple). A conserved sequence shared by four of the nudipleurans specifically, which show higher sequence similarity overall, is boxed (navy blue). **B.** The conserved boxed region in A is shown here as an 18 amino acid long sequence in the middle of the ELH gene. Shared among these four nudibranchs, it may be similar in function to additional, highly clade specific bioactive peptides in other gastropods that are shorter than ELH on the same gene. **C.** Multiple sequence alignment between the nudipleuran ELH sequences (top, purple), known mollusc ELH sequences (bottom, salmon), and an outgroup of DH44 from *Drosophila melanogaster.* Different colors around the letters denote different families of amino acids, indicating the level of conservation at that site. **D.** A maximum likelihood tree made from the multiple alignment in C to show the relationships between the short, bioactive peptide sequences (yellow segment in A). The Nudipleura forms a monophyletic clade. Numbers at nodes are bootstrap values, branch lengths are meaningful with tree scale at right. **E.** Multiple sequence alignment between the *B. stephanieae* ELH sequence and 4 other gastropods’ uncharacterized sequences on NCBI, that are likely also ELH orthologs lacking annotations, discovered with a Blastp search using the sequence of *B. stephanieae*.

Among the nudibranch samples with full-length sequences *(B. stephanieae*, *H. crassicornis*, *Armina spp*., and *F. iodinea*), which also cluster most closely in amino acid sequence similarity, there is an additional region of high amino acid similarity several hundred amino acids upstream of the terminal bioactive peptide (Figure 1B). At 18 amino acids in length, its strong conservation may indicate that it is an additional bioactive peptide, which may contribute to egg-laying behavior, similar to the alpha and beta bag cell peptides in the *Aplysia* genus (Nambu and Scheller, 1986). Its absence in *M. leonina* and *D. iris* may be due to the sequences being partial (e.g., lacking a signal peptide so lacking an unknown number of amino acids at the N-terminus). It also could be that their preprohormone is diverged significantly enough that the short bioactive peptides can longer be recognized and are also no longer bioactive across species, as is the case with the alpha and beta bag cell peptides of *A. californica* which are not active outside of the Aplysia genus. Indeed, it is also absent from the complete preprohormone of ELH in *P. californica*, which is the most distantly related nudipleuran that we examined.

The seven newly recovered bioactive nudipleuran putative ELH sequences were 37-40 amino acids in length, as expected, but differed from the previously known molluscan bioactive ELH sequences in several locations especially nearing the C-terminus as illustrated by a multiple sequence alignment (Figure 1C). Whereas other molluscs contain many arginine (R) and leucine (L) residues near the C-terminus, this shifted to asparagine (N) and methionine (M) in the Nudipleura. The nudipleurans also shared a conserved glutamine (Q) residue in the middle of the peptide at position 19, and nearly show consensus for an arginine (R) residue at position 20 (except for *M. leonina*) that the other molluscs have not conserved as strongly across the group. A maximum likelihood phylogenetic tree of this peptide revealed that the nudipleuran sequences were divergent enough from other molluscs that they cluster together monophyletically (Figure 1D).

To show the utility of the divergent nudipleuran ELH sequences, they were used to investigate published but uncharacterized gastropod protein sequences on NCBI using Blastp with *B. stephanieae*’s preprohormone sequence. Matches were found in the sacoglossan sea slugs *Elysia marginata* (accession GFS27520.1), *Elysia chlorotica* (accession RUS70186.1), *Plakobranchus ocellatus* (accession GFO26264.1), and the freshwater snail *Bulinus truncatus* (accession KAH9488389.1). Further examination of these uncharacterized genes reveals that the conserved bioactive region is 37-40 amino acids long and aligns very closely with *B. stephanieae*’s ELH sequence (Figure 1E). In all four of these species, this putatively bioactive peptide sequence occurs at the end of a longer gene, further evidence that these are likely to be the egg-laying hormone orthologs in these species.

### Synthetic ELH peptide induced egg-laying in *B. stephanieae*

Egg-laying behavior in *B. stephanieae* consists of continuous counterclockwise turning, from the inside and working outward, to produce a regularly spaced whorl of several hundred egg capsules (Figure 2A). Egg-laying behavior is accompanied by continuous rasping of the mouth, whether the animal was floating upside down on the water to lay eggs or attaching its egg mass to a surface.

**Figure 2:**
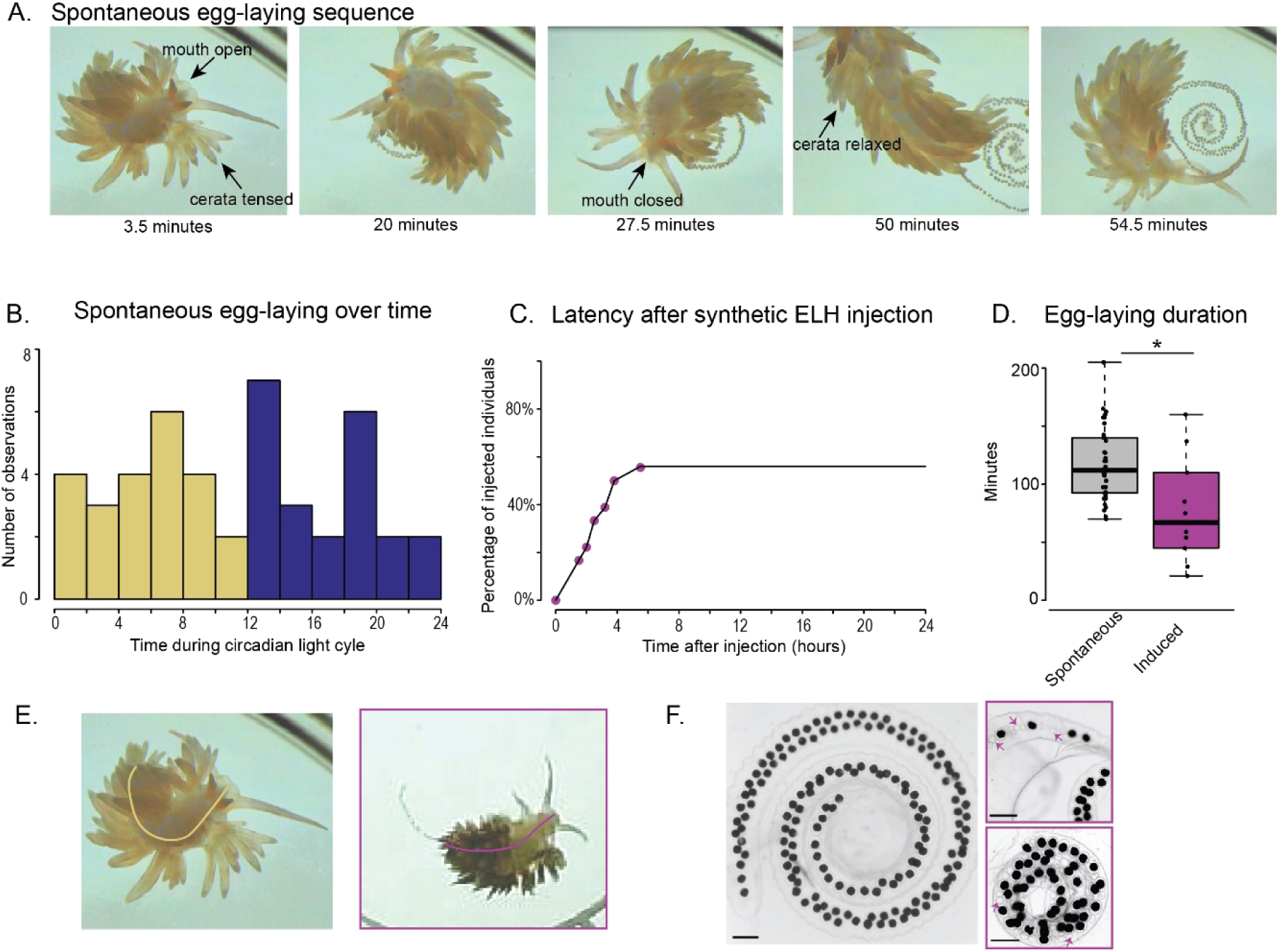
Spontaneous and induced egg-laying of the nudibranch *B. stephanieae*. **A.** Time sequence of spontaneous egg-laying postures in *B. stephanieae*. Note the tightly curled body, the “tension” in the cerata (arrows), and the changes in the mouth indicative of rasping (arrows). **B.** Histogram showing the number of spontaneous egg-laying events over a 24-hour standard day. 56 individual *B. stephanieae* were tracked over multiple days with Time=0 being 8am when the lights turned on. There was a total of 45 egg-laying events with no obvious temporal pattern. **C.** Cumulative-histogram of egg-laying events during the initial 24 hours after injection with synthetic ELH. All 10 egg-laying events (55% of 18 injected individuals) began within 6 hours of injection. **D.** Duration of egg-laying events compared between spontaneous (n=45) and induced (n=10) individuals. The difference was significant (Student’s T-test, p<0.05). **E.** When laying eggs spontaneously (left) individuals have markedly different behavior than when injected with ELH (right, magenta box) including a different body posture resulting in less turning and eggs laid in a strand rather than whorl (yellow curve, left; magenta curve, right). **F.** Egg quality differs between spontaneous (left) and induced (right, magenta boxes) egg-laying events. Notably, some egg capsules are also empty (magenta arrows). In all images, scale bars are 500 µm, illustrating how more eggs are laid spontaneously than when animals are induced to lay.

To determine the normal timing of egg-laying, 56 individuals were continuously video recorded for three- or five-day periods (see methods), for a total of 6144 “slug hours”. There was no temporal pattern to this population-level egg-laying; eggs were laid regularly across this entire period (Figure 2B). Egg laying was rare; only 45 events for the group were observed. There was only a 0.73% chance of an egg-laying event in any given hour. The average duration of such spontaneous egg-laying was 120 +/− 36 minutes (Figure 2D).

When synthesized *B. stephanieae* ELH peptide was injected into animals that were known to have previously laid eggs, 10 out of 18 individuals laid eggs within 6 hours (Figure 2C). The mean onset to egg-laying following the injection was 166 minutes. As a control, six animals were injected with saline; none of the controls laid eggs in the subsequent 24 hours. Therefore, ELH injection had a significant impact on the proportion of slugs laying eggs versus injection of saline (2-tailed Fisher’s Exact Test, p = 0.046). Within those 6 hours, there was a 9.25% chance of observing an egg-laying in any given hour, an increase over spontaneous egg-laying.

Artificially induced egg-laying differed from spontaneous egg-laying in several ways. The average duration of egg-laying in injected animals was 77.5 +/− 44 minutes (N=10), which was significantly shorter than the 120 minutes seen in spontaneous events (Student’s T-test, p<0.05, Figure 2D). Furthermore, individuals that had been injected with ELH showed less of the stereotypical curl of their body and less turning (Figure 2E). Moreover, the egg masses that were laid after ELH injection were not as uniform as spontaneously laid egg masses. Seven of the ten egg masses showed some abnormality, such as the sleeve being empty of egg capsules entirely or major structural differences in the whorl shape of typical egg masses as shown in Figure 2F. After two weeks, four of the seven abnormal egg masses produced viable embryos, whereas the three normal egg masses all contained viable embryos.

### ELH was expressed in identifiable neurons in all ganglia

Neurons in each ganglion of the *B. stephanieae* brain were labeled by HCR probes specific for ELH preprohormone (Figure 3, Figure 4A, Table 1). Many of the neurons were recognizable by their soma position and size across individuals at the same stage of development as well as across different stages of post-metamorphic development. We numbered the neurons and clusters of neurons that were consistently present in the adult and put them in an order that indicated when they were recognizable in juvenile stages (Figure 3). There were 18 consistently identifiable classes of ELH-expressing single neurons or neuronal clusters and three variably present classes (Figure 3, Table 1). Although the buccal ganglion (*bcg*) has ELH-expressing neurons (Figure 4B), it was not included in this list because its cylindrical shape and long connective made it difficult to orient consistently, adding to the uncertainty of identifying neurons.

**Figure 3:**
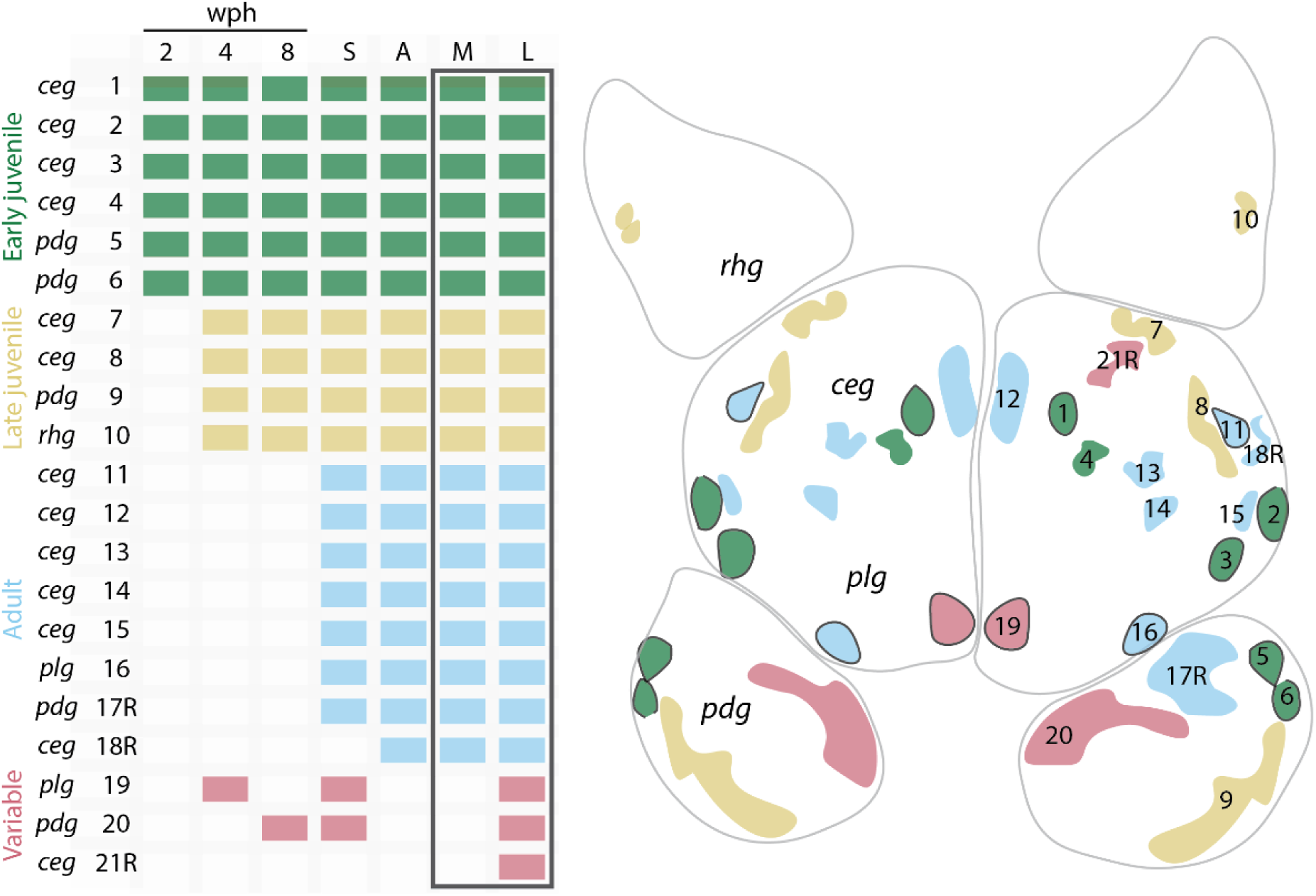
Characteristics of the mapped ELH-positive neurons across developmental stages and reproductive states in the *B. stephanieae* brain. **A.** ID numbers 1-18R correspond with the classes described in Table 1, IDs 19-21R were not seen in the standard brain but are seen in selected stages/states. ID numbers with “R” indicate clusters present only on the right. The columns represent a loose developmental continuum with neuron types arranged in order of appearance and colored corresponding to four types based on their appearance in the brain: “early juvenile”, “late juvenile”, “adult”, and “variable”. The adult brain samples at different reproductive states are boxed in black. **B.** Schematic of the brain showing the approximate locations of all 21 ELH-expressing neurons. Only the neurons on the right half are labeled. Abbreviations - wph: weeks post-hatching, S: sub-adult, A: adult, M: mating, L: egg-laying, *rhg*: rhinophore ganglion, *ceg*: cerebral ganglion, *plg*: pleural ganglion, *pdg*: pedal ganglion

**Figure 4:**
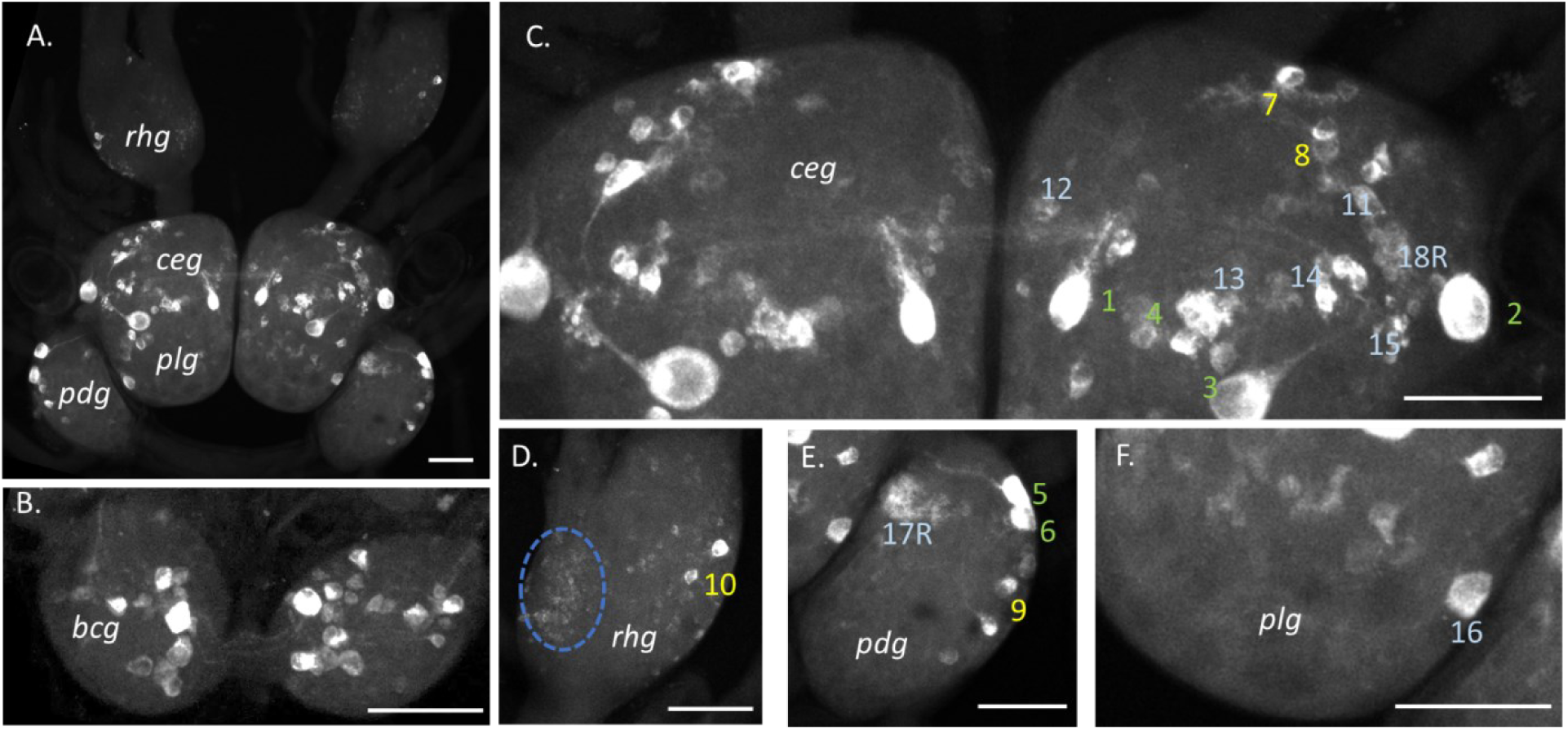
*In-situ* HCR fluorescent labeling of ELH mRNA in neurons of adult *B. stephanieae.* Neurons showing ELH expression in the whole brain (**A**), buccal ganglion (**B**, *bcg*), cerebral ganglion (**C**, *ceg*), rhinophore ganglion (**D**, *rhg*), pedal ganglion (**E**, *pdg*), and pleural ganglion (**F**, *plg*). The *rhg* contains dozens of small neurons (**D**, dotted blue circle) and only two identifiable neurons (Cluster 10). In all panels, individual neurons or clusters of neurons that seen in >50% of cases are numbered as in Table 1. In all images, scale bars are 100 µm.

**Table 1.**
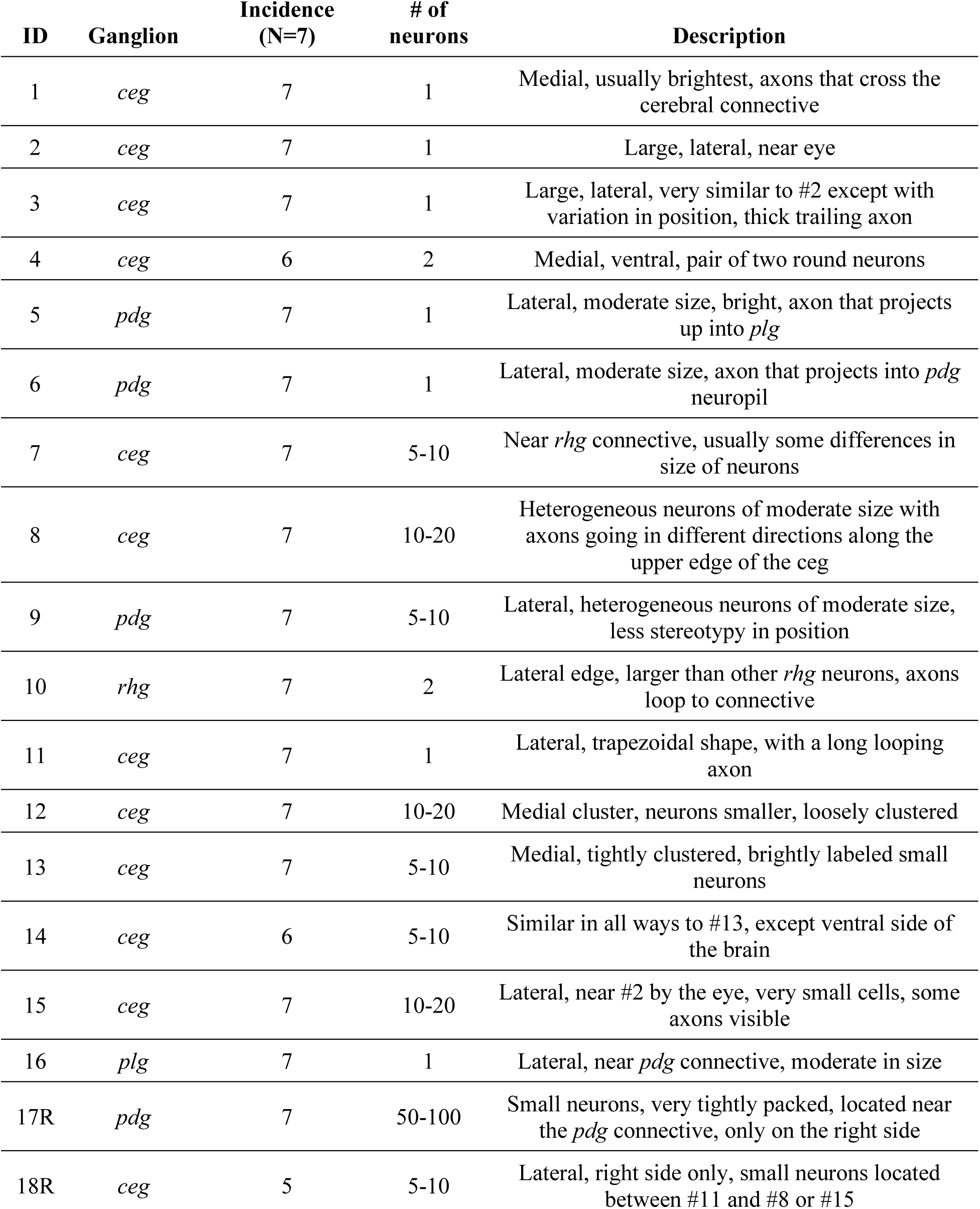
Characteristics of the mapped ELH-positive neurons in a standard *B. stephanieae* brain. Identification numbers that have an “R” indicates presence only on the right side. The number of observations under each category is reported. Abbreviations: *ceg* - cerebral ganglion, *plg* – pleural ganglion, *pdg*-pedal ganglion, *rhg* – rhinophore ganglion.

| ID | Ganglion | Incidence<br>(N=7) | # of<br>neurons | Description |
| --- | --- | --- | --- | --- |
| 1 | *ceg* | 7 | 1 | Medial, usually brightest, axons that cross the cerebral connective |
| 2 | *ceg* | 7 | 1 | Large, lateral, near eye |
| 3 | *ceg* | 7 | 1 | Large, lateral, very similar to #2 except with variation in position, thick trailing axon |
| 4 | *ceg* | 6 | 2 | Medial, ventral, pair of two round neurons |
| 5 | *pdg* | 7 | 1 | Lateral, moderate size, bright, axon that projects up into *plg* |
| 6 | *pdg* | 7 | 1 | Lateral, moderate size, axon that projects into *pdg* neuropil |
| 7 | *ceg* | 7 | 5-10 | Near *rhg* connective, usually some differences in size of neurons |
| 8 | *ceg* | 7 | 10-20 | Heterogeneous neurons of moderate size with axons going in different directions along the upper edge of the *ceg* |
| 9 | *pdg* | 7 | 5-10 | Lateral, heterogeneous neurons of moderate size, less stereotypy in position |
| 10 | *rhg* | 7 | 2 | Lateral edge, larger than other *rhg* neurons, axons loop to connective |
| 11 | *ceg* | 7 | 1 | Lateral, trapezoidal shape, with a long looping axon |
| 12 | *ceg* | 7 | 10-20 | Medial cluster, neurons smaller, loosely clustered |
| 13 | *ceg* | 7 | 5-10 | Medial, tightly clustered, brightly labeled small neurons |
| 14 | *ceg* | 6 | 5-10 | Similar in all ways to #13, except ventral side of the brain |
| 15 | *ceg* | 7 | 10-20 | Lateral, near #2 by the eye, very small cells, some axons visible |
| 16 | *plg* | 7 | 1 | Lateral, near *pdg* connective, moderate in size |
| 17R | *pdg* | 7 | 50-100 | Small neurons, very tightly packed, located near the *pdg* connective, only on the right side |
| 18R | *ceg* | 5 | 5-10 | Lateral, right side only, small neurons located between #11 and #8 or #15 |

The cerebral ganglion (*ceg*, Figure 4C) and the rhinophore ganglion (*rhg*, Figure 4D) had the largest numbers of ELH-expressing neurons and the pleural ganglion (*plg*, Figure 4F) had the fewest. In larger neurons within most ganglia, neurites were discernable close to the somata, indicating the presence of mRNA there and translation potentially occurring outside of the cell body.

The ELH expressing neurons in the *ceg* varied greatly in size and labeling intensity within animals. However, their morphologies and locations were stereotyped across individuals (Figure 4C), allowing 12 classes to be reliably identified (Table 1). Several prominent neurons had axons consistently visible, with their projection patterns aiding in their identification (Figure 4C, Neurons 1 and 2). Some neurons were more variable in position (Figure 4C, Neuron 3). One ELH-expressing neuron had a unique trapezoidal shape and characteristic looping axon (Figure 4C, Neuron 11). Several smaller neurons could not be differentiated as individuals but were consistently clustered together in stereotyped locations (Figure 4C, Clusters 13 and 14). Some clusters were more diffuse (Figure 4C, Clusters 7 and 12). A specific cluster of small neurons was present only on the right side of the *ceg* (Figure 4C, Cluster 18R).

In the lateral region of each *rhg*, two neurons consistently expressed ELH (Table 1, Figure 4D). Usually, a neurite was present emerging from each of the two neurons, which looped up then down towards the rhinophore connective. These neurons were noticeably larger than the rest of the very small (5-10 micron), numerous ELH-positive neurons spread diffusely throughout the rhinophore ganglion, which were not further catalogued here.

Neurons in the *pdg* (Figure 4E, Table 1) were concentrated along the lateral edge of the ganglion and were generally more consistent in their location than neurons in the *ceg*. Some could be identified as individuals by the visible segments of their axons (Table 1, Figure 4E, Neuron 5 and 6). One cluster was found only in the right *pdg* and consisted of 50-100 small neurons near the pleural-pedal connective (Table 1, Figure 4E, Cluster 17R). Neurons in the *plg* (Figure 4F, Table 1) were less intensely labeled and more variable in position than the other ganglia. Only one was consistently identifiable, located along the posterior edge of the ganglion, bordering the *pdg* (Figure 4F, Table 1, Neuron 16).

### ELH expressing neurons appeared at different times during juvenile development

ELH-expressing neurons that were identified in the adult could be recognized in post-metamorphic juveniles 2, 4, and 8 weeks post-hatching. The brains at these stages were too small for dissection and were thus labeled in-place within the body (Figure 5A). The positioning of the ganglia in the body of the juvenile differed from how the dissected brain laid on a glass slide, such that the *rhg*, *ceg*, and *plg* are dorsal and the *pdg* and *bcg* sat ventrally, necessitating separate imaging from the dorsal (Figure 5Bi, Ci, Di) and ventral (Figure 5Bii, Cii, Dii) orientations.

**Figure 5:**
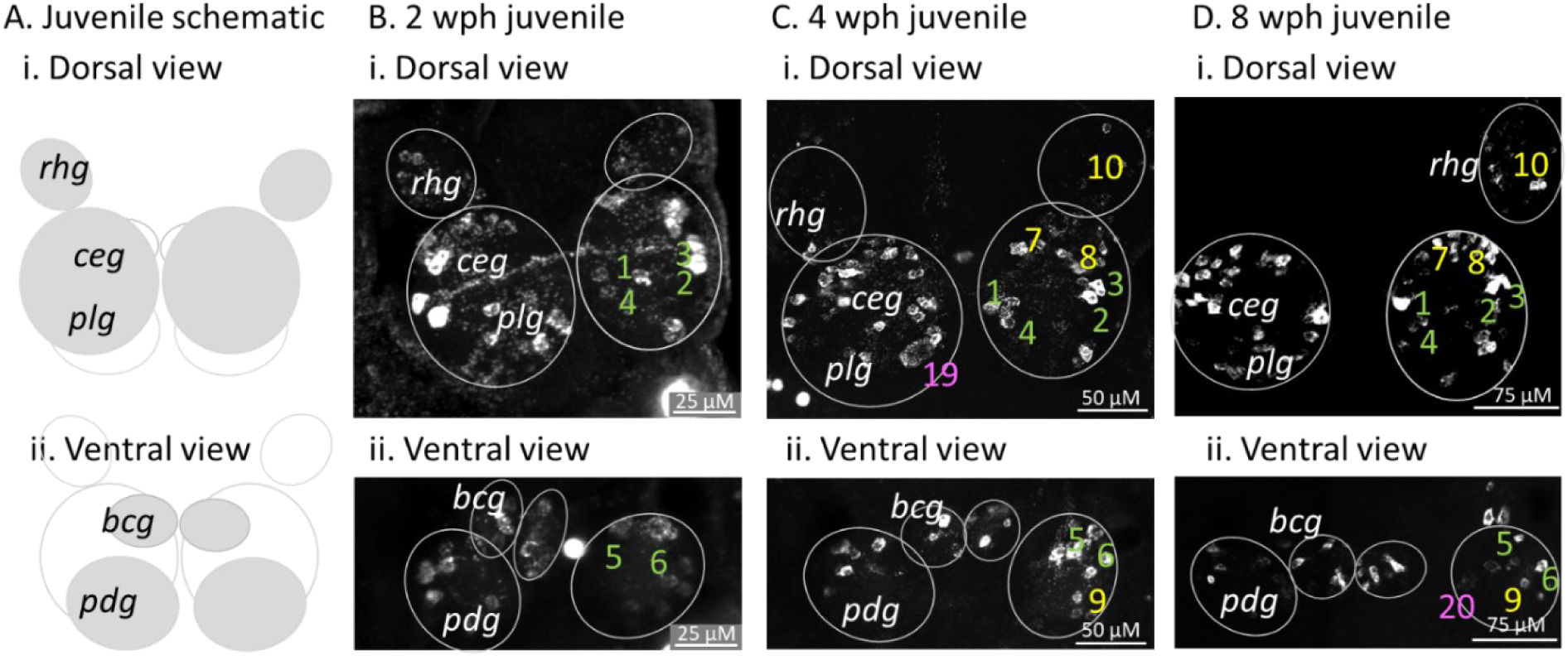
ELH expression in the brains of early, middle, and late stage juveniles. **A.** Dorsal (i) and ventral (ii) schematics of the juvenile *B. stephanieae* brain as viewed in place as a wholemount, with visible ganglia on the dorsal versus ventral side as solid circles. **B.** Early stage, 2-week post hatching juveniles already have ELH-expressing neurons that correspond to those in the adult in the *ceg*, neurons 1-4 (i) and the *pdg*, neurons 5-6 (ii). **C.** Middle stage, 4-week post hatching juvenile brains increased the numbers of neurons and clusters corresponding to those seen in adults in the *ceg,* neurons 1-4 and 7-8, with cluster 10 also in the *rhg* (i). On the left side, neuron 19 was found in the *plg*, which was not seen in the standard adult brain. In the *pdg* were all of the same neurons as in adults, neurons 5-6 and 9 (ii). **D.** In an 8-week post-hatching late juvenile, no identifiable new neuron types were added although significant growth of the ganglion occurred (i). Within the *pdg*, Cluster 20 is seen, which was not in standard adults (ii). Asymmetric Cluster 17R, which is seen in adults, is not yet present in late juveniles at 8 weeks post-hatching. Note that the buccal mass, which abuts the *bcg*, is incompressible and often deforms the position of the ganglia when mounting and causing the two *pdg* to vary in position relative to each other and other ganglia.

Despite these challenges, it was possible to locate several classes of adult neurons even in the earliest juvenile stages. In the dorsal view, in the *ceg* (Figure 5 Bi, Ci, Di), neurons that were individually identifiable (Neurons 1, 2, 3, and 4) were consistently present through all juvenile stages. Neurons that occurred in clusters were only recognizable in the *ceg* later in development (Clusters 7 and 8, Figure 5Ci, Di). The two larger neurons of the adult *rhg* were also not seen until later in development (Cluster 3, Figure 5Ci, Di). In the ventral view, the *pdg* and *bcg* also had numerous ELH-expressing neurons that were identifiable during the earliest juvenile stages; these identifiable neurons were seen through the majority of juvenile stages, especially in the *pdg* (Figure 5 Bii, Cii, Dii, Neurons 5, 6, and Cluster 9).

Across all juvenile stages, no right-sided asymmetric neuron clusters were seen (Clusters 17R and 18R). In some animals in the mid and late juvenile stages, neurons that were not seen in the standard adult brain were observed, specifically one in the *plg* (Figure 5 Ci, Neuron 19) and a cluster in the *pdg* (Figure 5 Di, Cluster 20). These had dimmer labeling that was punctate, indicating lower expression.

### ELH expression varied across reproductive states in specific neurons

We suspected that the appearance of neuron types expressing ELH at low levels in mid- and late-stage juveniles (Figure 5, neuron 19 and cluster 20), which was not seen in the standard adult, was related to reproduction. In the present study, sub-adults were communally housed and although they had not laid eggs yet and had no eggs visible in their reproductive organs when they were sacrificed, these animals likely had already mated. Precocious mating with early sperm donation, weeks prior to egg-laying, was recently reported in *B. stephanieae* (Taraporevala et al., 2022). We found that in sub-adults capable of this behavior (N=5), all 18 neuron classes seen in adults were identifiable (Figure 6A). This included clusters that were asymmetric on the right side. It also included neurons in the *plg* and *pdg* (Figure 6A, neuron 19 and cluster 20), which had been sporadically noted in younger juvenile stages, and a cluster in the right *ceg* (21R). These neuron classes, not seen in standard adult brains, were again noted to show more punctate labeling indicating lower expression. The presence of these neurons versus the standard brain were particularly obvious because of where they appear: medially in the *plg* (Figure 6A, neuron 19) and medially in the *pdg* (Figure 5A, Cluster 20), which are distinct from any regularly occurring neurons.

**Figure 6:**
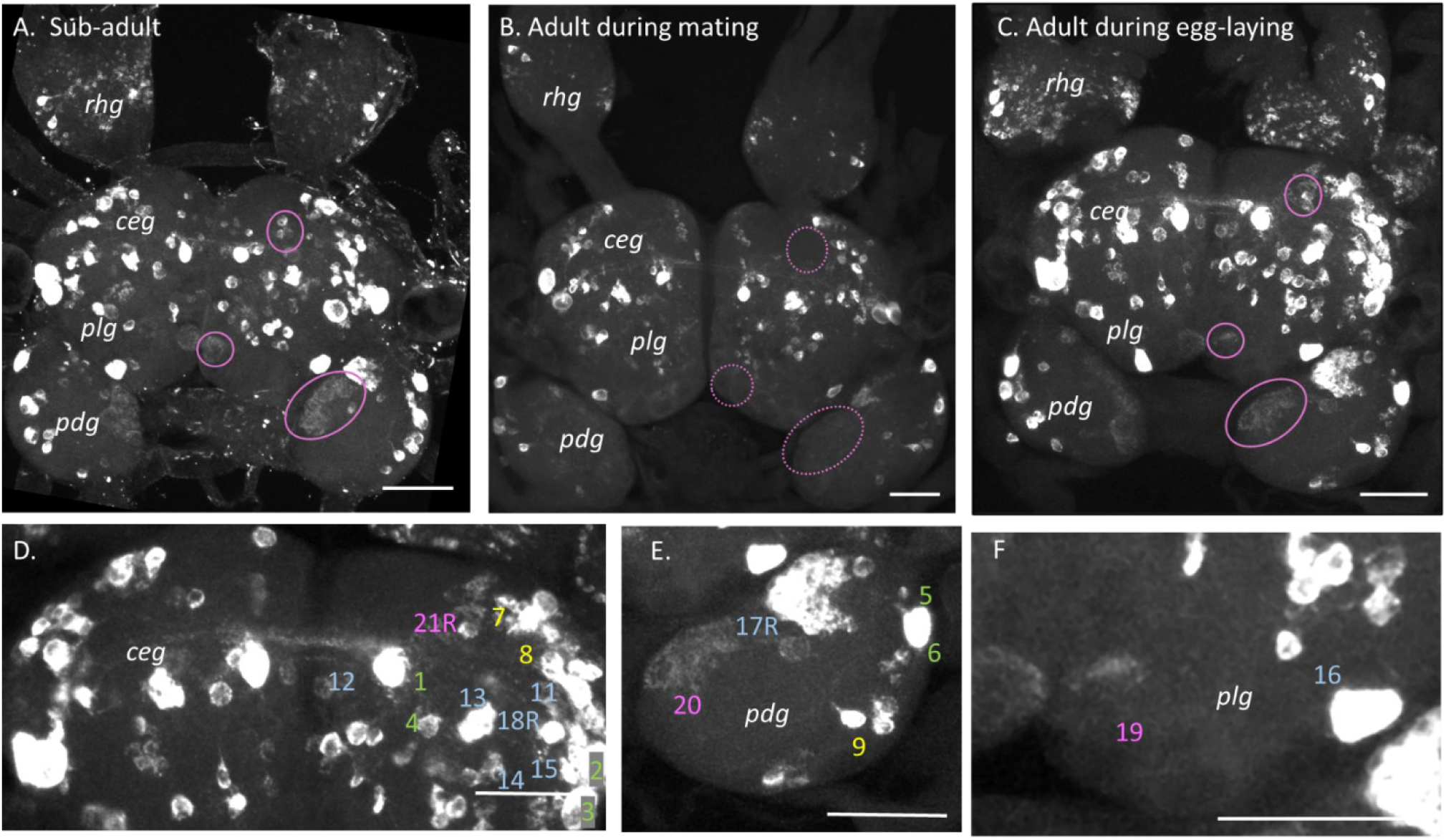
ELH in *B. stephanieae* across different reproductive states highlights expression in three neuron classes. View of the cerebral (*ceg*), pleural (*plg*), pedal (*pdg*), and rhinophore (*rhg*) ganglia showing ELH expressing neurons. **A.** Sub-adult brain labeled for ELH before it had ever laid eggs but after it had free access to mates for several weeks. Neurons not seen in a normal adult brain are circled in pink (n=5). **B.** Brain of adult sacrificed during the first hour of mating with dotted pink circles showing where the same neurons in A and C are not present (n=4). **C.** Brain of adult sacrificed during the first hour of egg-laying with pink circles showing neurons seen in sub-adults in A but not seen in a normal adult brain (n=5). **D-F.** Higher magnification views of *ceg* (D), *pdg* (E), and *plg* (F) of the egg-laying adult in C. Identified neurons and clusters are labeled, including asymmetrical Cluster 21R, that occurs only in the ventral anterior right *ceg* (D), bilateral Cluster 20, in the *pdg* (E), large Neuron 19 on the medial posterior edge of the *plg* (F). All scale bars are equal to 100 µm.

To determine whether the presence of specific neurons was related to reproductive behavior, individuals that were in the act of mating were sacrificed. These animals were paired together for at least 1hour but no more than 3 hours (n=4). During that time, animals were anaesthetized, and the anterior half of their body was severed and fixed, with the brain dissected free of the tissue only after fixation. The primary 18 classes of neurons were labeled (Figure 6B). Of the neurons described in the pre-reproductive late-stage juveniles and sub-adults (IDs 19-21R), none were present in more than 50% of observed samples.

Finally, we labeled ELH expression in the brains of individuals that had been laying eggs. Animals were observed laying eggs for at least 20 but less than 60 minutes. They were then anaesthetized, fixed, and later dissected (Figure 6C, n=5). All 18 classes of ELH-expressing neurons described in non-egg-laying adults were observed in egg-laying individuals. There were, however, again the additional neuron types that had been seen in some juveniles and sub-adults, now occurring in the animals engaged in laying eggs. Cluster 21R, was seen in the right *ceg* in 3 out of 5 egg laying individuals (Figure 6D). Cluster 20, consisting of 4-5 large neurons, appeared on the medial edge of each *pdg* in 4 out of 5 of egg-laying adults, but in none of the non-egg-laying adults (Figure 6E). In 3 out of 5 individuals laying eggs, one very large *plg* Neuron 19, was present medially in both the right and left ganglion (Figure 6F).

### ELH-expressing cells are located in peripheral sensory tissues

*B. stephanieae* uses paired oral tentacles, rhinophores, and its mouth for chemosensory tasks. Numerous small cells expressing ELH were found in these peripheral structures. HCR labeling within cells was punctate, indicating lower expression than in the ganglionic neurons. These cells were most dense in the tissues surrounding the mouth, especially the lower lip (Figure 7A). This population of cells were contiguous with those at the base of the oral tentacle (Figure 7B) and appear along the full length of the tentacles to their tapering tips (Figure 7C). The ELH-positive cells along the mouth and oral tentacles were located near the surface, in the sensory epithelium. Many of these cells had neurites projecting through the epithelium to the surface (Figure 7A-C, insets) suggesting they are sensory neurons.

**Figure 7:**
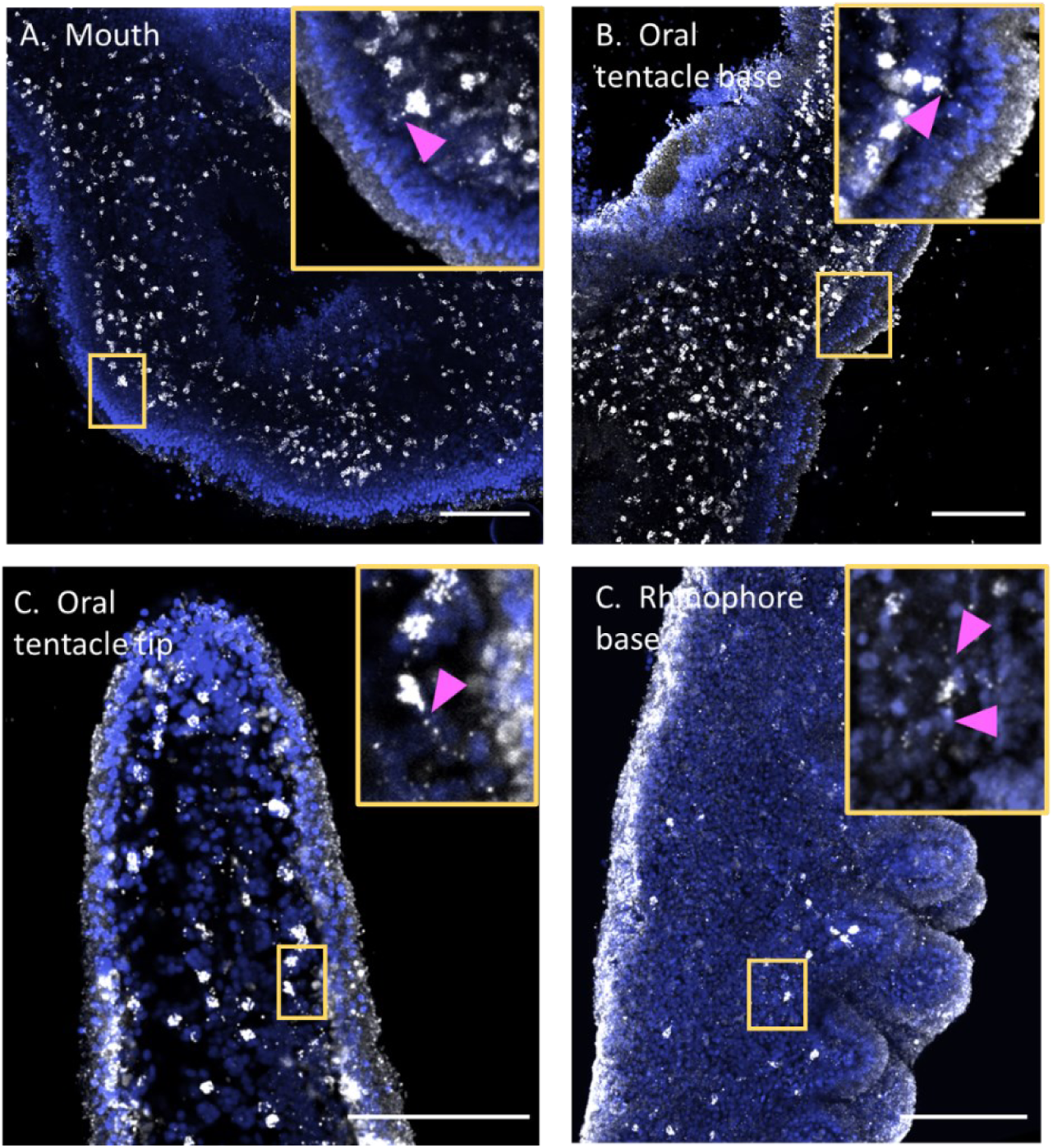
ELH expression in the sensory appendages. **A.** ELH-expressing cells (white) are spread densely across the mouth and lips, in the sensory epithelium. DAPI (blue) is used as a co-stain for cell nuclei, to show the shapes of the appendages. **B.** ELH-expressing cells on the ventral surface of the oral tentacles, appearing continuous with those on the lips surrounding the mouth (bottom left corner). **C.** A close-up of ELH-expressing cells on the tip of the oral tentacle. **D.** The rhinophore is covered sparsely with a population of cells that expresses ELH that occurs deep underneath the epithelial layer. In all panels, yellow-boxed insets show cells which are putatively neurons with visible neurites projecting out through the epithelium (A, B, C, magenta arrowheads), indicating they’re sensory neurons, or neurites projecting in both directions (D, magenta arrowheads). All scale bars (bottom right) are 100 µm.

ELH-expressing cells were present sparsely throughout the length of the rhinophores (Figure 7D). These were located deeper in the tissue. Some of them were bipolar neurons with visible neurites (Figure 7D inset); their location deep in the appendage may indicate they are a type of interneuron.

## Discussion

We found orthologous ELH preprohormones in several nudipleuran molluscs, with the predicted bioactive peptide showing clade-specific sequence differences. To verify behavioral activity, the short bioactive peptide ELH from *B. stephanieae* was synthesized and injected into *B. stephanieae*. *In-situ* HCR labeling showed consistent ELH-expressing neurons throughout the brain of adult animals. These identifiable neurons were seen to be sequentially added over the course of juvenile development. Certain neurons exhibited ELH-expression only in late-stage juveniles and animals that were actively laying eggs. Appearance of neurons in later stages of juvenile development may imply a role in reproductive physiology and behavior that only occurs at sexual maturity. One such group of neurons, an asymmetric cluster in the right pedal ganglion (cluster 17R), may be homologous to the main ELH secreting bag cells of *Aplysia californica* and CDCs of *Lymnaea stagnalis.* It may also be possible that late-developing cerebral neurons in *B. stephanieae* (clusters 18R and 21R) may be homologous to those seen in *A. californica* which have descending control over other clusters (Brown et al., 1989). ELH was also consistently expressed by putative peripheral sensory neurons. Such widespread expression had not been described in other molluscs but implicates ELH has having pleiotropic function - as a general signaling molecule, impacting *B. stephanieae*’s physiology beyond reproduction.

### Sequence differences in the ELH ortholog in the Nudipleura

The predicted amino acid sequences of the ELH peptide orthologs in the Nudipleura have notable differences at their C-terminus, which may explain why ELH had not been previously described in this clade. Behavioral validation with a synthesized peptide was key to show its homology to ELH in other gastropods. Differences in the C-terminus of the peptides suggest that the receptor for this peptide may have diverged significantly in this group and, if so, it would be interesting to determine which parts of the receptor sequence have changed.

Differences in sequence near the C-terminus of the nudipleuran ELH genes were extensive enough that we used both Blastp and HMMER to be confident that we had identified proper sequences. This underscores the importance of manual investigation in discovering conserved neuropeptides. For example, a quick Blastp search on NCBI with the amino acid sequence of *B. stephanieae*’s ELH preprohormone returned good matches to hypothetical proteins of the heterobranchs *Elysia marginata*, *Elysia chlorotica*, *Plakobranchus ocellatus*, and *Bulinus truncatus* (Figure 1E). This shows that divergence in the ELH peptide sequence, as found by studying nudipleurans, may extend to other heterobranch gastropods as well. The use of manual, iterative investigation will aid in annotation of genes previously listed as hypothetical or uncharacterized.

### Induced egg-laying behavior in *B. stephanieae*

Egg-laying in response to injection of synthetic ELH differed from spontaneous egg-laying in timing, duration, egg quality, and turning behavior. These behavioral characteristics are likely linked in spontaneous egg-laying: *B. stephanieae* locomotes at specific speed and bends at a specific angle to produce a consistently “packed” whorl of eggs and continues as long it has eggs to deposit. Injection of ELH may hijack this sequence of events and result in inconsistencies compared with the spontaneous egg-laying process. Additionally, we injected only the bioactive neuropeptide, 37-40 amino acids at the terminus of a much longer gene. It seems likely that there are shorter neuropeptides located on the same gene (e.g., like the alpha and beta bag cell peptides in *Aplysia spp.*). These were absent in our assay but could modulate specific aspects of egg-laying behavior in *B. stephanieae*.

The effects of ELH injections in other gastropods produce behavioral phenotypes like what we have described. In the pond snail, *Lymnaea stagnalis,* the differences between spontaneous and induced-via-injection egg-laying behavior have been carefully analyzed. Injected *L. stagnalis* go immediately into a turning phase, without a quiescent period, resulting in less uniform egg-masses (Ter Maat et al., 1989). In the garden snail, *Theba pisana*, digging and turning behavior to deposit egg masses was sometimes seen upon specific dosages of injected, synthetic ELH, without any eggs being laid (Stewart et al., 2016).

### Distribution of neurons expressing ELH in *B. stephanieae*

*Berghia stephanieae* has more neurons in its brain overall that express ELH than was previously reported for either *A. californica* or *L. stagnalis*. *A. californica* was reported to have only 20 neurons outside of the bag cell cluster (McAllister et al., 1983) and in *L. stagnalis* there are over 100 other than the CDCs (van Minnen et al., 1988). However, *B. stephanieae* has hundreds in the rhinophore ganglia alone, with an estimated total of 300-400 neurons brain wide.

Our developmental series makes it possible to pinpoint neurons that develop in the post-hatching period. Studies in *L. stagnalis* have found that amount of this hormone steadily increases as individuals grow and develop (Dogterom et al., 1983), and that the neurons that produce this hormone, the CDCs are specifically not present until later in development (Roubos et al., 1988). In *B. stephanieae*, we have seen that specific classes of ELH-expressing neurons do not appear until the sub-adult stage of development. Neurons that only appear in stages shortly before sexual maturity may play more of a direct role in egg-laying; this includes the CDCs and bag cells in other taxa. Namely, the cluster of up to 100 small neurons in the right pedal ganglion of *B. stephanieae* also appeared only beyond the 8 week post hatching phase, at some point before the sub-adult stage (Cluster 17R). They are asymmetric only on the right side, and in both *A. californica* and *L. stagnalis* there are more neurons in the right clusters of the bag cells or CDCs, which correspond to their lateralized, right-sided reproductive organs. This, combined with location: near the pedal connective to the cerebropleural ganglion, a potential neurohemal release zone, makes it possible that they are directly involved in egg-laying behavior. A manipulative study on *Pleurobranchaea californica* suggests both of the pedal ganglia as sites for the bag cell homologous neurons (Ram et al., 1977).

However, there were also other clusters in the *ceg* of *B. stephanieae*, some of which are also right-sided asymmetrical (e.g., Cluster 18R) which do not appear until sub-adult stages (Table 1). These clusters are on the dorsal-most and ventral-most faces of the ganglion, which could aid in release. These clusters could be linked in a circuit of descending control as speculated in *A. californica* (Brown et al., 1989).

Some of the other neurons expressing ELH in *B. stephanieae* were present in the earliest juvenile stages. These neurons can be correlated with neurons seen in adult *B. stephanieae*. They are individually identifiable by morphology and ELH-expression across developmental and reproductive stages (post-mating sub-adults, mating adults, egg-laying adults, and standard adults). They could be divided into specific classes. This, combined with the fact that a vast number of sensory neurons express ELH, implies that ELH is a key general signaling molecule in *B. stephanieae*.

### Upregulation of ELH expression in specific neurons

We found that specific neurons consistently express ELH during active egg-laying but not in standard “at rest” adult nudibranchs. These neurons occur in the *ceg*, *pdg*, and *plg*. This is congruent with findings in *P. californica ceg*, where injections with homogenized extracts of *ceg* only induced egg-laying if they were made from individuals sacrificed during active egg-laying (Ram et al., 1977). Similarly, in *L. stagnalis*, it was the pedal ganglia that have been recognized as essential for generating the turning behavior required for egg-laying (Hermann et al., 1994). However in the *pdg* of *L. stagnalis* ELH expression was only noted in neuronal fibers and not somata (van Minnen et al., 1988). *B. stephanieae* has several ELH expressing classes in the *pdg*, so this may be a clade-specific difference in neuron location between these gastropods.

ELH that was visualized during egg-laying but not during mating suggests that the increased expression in these neurons is specific to egg-laying and does not occur all reproductive behaviors. Expression in these same classes of neurons in some sub-adults and late juveniles further suggests that there may be a build-up of ELH in preparation for the animal’s first egg-laying event. This is reminiscent of what was observed in *L. stagnalis*; transcription of ELH in young animals builds until it is released by translation which happens at the time of egg-laying (De Lange et al., 1994). Behavioral work in *B. stephanieae* recently showed that even virgin *B. stephanieae* that were isolated their entire lives eventually laid eggs, albeit weeks later than animals in their cohort that had access to mates (Taraporevala et al., 2022).

### Neuropeptides in the periphery

The extent of ELH expression in all peripheral tissues was unexpected. It was previously concluded in *A. californica* that ELH-expressing cells at the periphery migrate into the brain during development (McAllister et al., 1983). Our findings show that there is extensive expression of ELH in the periphery even in mature individuals, and that at least some subset of the peripheral cells expressing ELH have dendritic projections through the epithelium, meaning that they are likely to be mature sensory neurons.

In invertebrates, it is increasingly being recognized that neuropeptides are expressed in peripheral sensory neurons (Leinwand and Chalasani, 2011; Taghert and Nitabach, 2012). In *Octopus vulgaris*, a male-associated reproductive hormone, APGWamide, is expressed in sensory neurons of the olfactory organ (Polese et al., 2015), implying that the neuropeptides with reproductive function can modulate sensory neuron dynamics in a mollusc. This could be the case for ELH in *B. stephanieae*’s sensory appendages, particularly on the mouth and oral tentacles where ELH-expressing cells are much more abundant. *B. stephanieae* does share an “inspection” phase with *L. stagnalis* during the egg-laying process (Ter Maat et al., 1989): they touch their egg mass after it is deposited on the substrate. *B. stephanieae* was observed to do this with their oral tentacles and mouth (Figure 2A).

## Concluding remarks

The consistent expression of ELH in many neurons, some of which are identifiable, in adult and juvenile *B. stephanieae* seems to imply function beyond egg-laying. Indeed, ELH has been recently placed within the corticotropin-releasing hormone gene family (Mirabeau and Joly, 2013), which is involved broadly with stress response and known to have expanded into many more specialized roles in vertebrates (Lovejoy and Hogg, 2021). Additional studies are needed to determine the functional roles played by the diverse sets of neurons in *B. stephanieae*.

## Acknowledgements

We would like to first acknowledge Dr. Adriano Senatore for generation of the unpublished nudipleuran brain transcriptomes that we assembled and analyzed in this study. We would also like to thank Phoenix Quinlan and Thi Bui for use of their modified IR-capturing webcam and IR LED floodlight, as well as Kristina Nedeljkovic for early troubleshooting of the ELH HCR protocol. Many thanks to Liz Cowley for her work on the graphical abstract. We also acknowledge the use of the confocal scanning microscopes and support from the Light Microscopy Facility and Nikon Center of Excellence at the Institute for Applied Life Sciences, UMass Amherst. The work included in this study was funded by the following grants: NIH U01-NS108637, NIH U01NS123972, NIH R01NS133654, and NSF IOS 2227963.

## Competing Interests

The authors declare no competing interests.

## Data Accessibility

Raw data will be made available upon request.

## Notes

### Competing Interest Statement

The authors have declared no competing interest.

