## Supplemental Table 1 for "Egg-laying hormone expression in identified neurons across developmental stages and reproductive states of the nudibranch *Berghia stephanieae*"

### Supplemental materials (12/21/2023)

| Mollusc species | Mollusc group | Source of ELH sequence for hmmer profile |
|---|---|---|
| *Lottia gigantea* | Gastropod | Veenstra (2010) |
| *Lymnaea stagnalis* | Gastropod | Vreugdenhil et al. (1985) |
| *Aplysia californica* | Gastropod | Scheller et al. (1983) |
| *Aplysia dactylomela* | Gastropod | Cummins et al. (2010) |
| *Aplysia parvula* | Gastropod | Nambu and Scheller (1986) |
| *Charonia tritonis* | Gastropod | Bose et al. (2017) |
| *Theba pisana* | Gastropod | Stewart et al. (2016) |
| *Mizuhopecten yessoensis (Patinopectin yessoensis)* | Bivalve | Zhang et al. (2018) |
| *Pinctada fucata* | Bivalve | Stewart et al. (2014) |
| *Crassostrea gigas* | Bivalve | Stewart et al. (2014) |

**Supplemental Table 1:** References for molluscan ELH sequences used to build hmmer profile and in blast

| Nudipleuran species | blastP e-value with A. californica ELH as query | blastP e-value with Dendronotus ELH as query | blastP e-value with Berghia ELH as query | HMMER e-value using molluscan ELH profile |
|---|---|---|---|---|
| *Berghia* | Not found | 0.003 | --- | 0.09 |
| *Dendronotus* | 0.21 | --- | 0.034 | 3.9e-09 |
| *Melibe* | 0.011 | 6e-11 | 0.096 | 5e-09 |
| *Flabellinopsis* | Not found | 0.14 | 5e-11 | 3.1e-04 |
| *Hermissenda* | Not found | Not found | 1e-08 | 0.75 |
| *Pleurobranchaea* | Not found | .017 | 2e-06 | 0.016 |
| *Armina* | Not found | Not found | 1e-09 | |

**Supplemental Table 2:** E-values to summarize results of blastP and hmmer searches for Nudipleuran ELH sequences

| Nudipleuran species | Data on annotated features | ELH preprohormone AA sequence |
|---|---|---|
| *Berghia* | Length: 374<br>Signal pep: 1-28<br>Cleavage sites: 11<br>Locations: 31, 39, 79, 148, 198, 262, 276, 308, 317, 333, 334, | MKTPLTNFIFDLAIMTTTMVAIISPVHATLRSRDLFADKSS PLPNLISTIKSLRLMNTCRQHVGQWKYNPNPSLYFPSRTF QNVNQNSIREMAKTTSYNDANFASKLINVIKNSKSSTSN LPLFQDASFLDNISVDLRKMLNEKLNKKFSLYSHSTKTD LNEQSNKNSFEFLEQMCLKYAFDKLLQNQNREQYLAGK KALEDRNHIVRNLQGLQQFPEMLSPETSGNVGLLSLLSP DVNIREFRHSFHMSTPPNHPVYNDGRSFSNTYNSNQKM ERSPFDASEGNIVPSSLAQIDPEVQQQDRSLAPKRSKLMA PSKRGFPSLFTWLGNFRNKRRHRLSVNNAMMTLSSMV MEDARQRAQKNRNNIKYHMYLQG |
| *Dendronotus* | Length: 267<br>Signal pep: N/A<br>Cleavage sites: 6<br>Locations: 22, 81, 106, 182, 196, 227 | MRTSLDQRRQRQRRRRRQRQRRSNDNGGLGATSYLSKL LYGGDNALTSQGEYQVEREPERPSSSSSSHVTTAILRMRR HHRASPARTRNIEMRYLCRKYALESLLRSDSYEDDDGYS ISHQLLNAKVYHSPNQYQTPSDERNIEQQQDQDTAASKA LLLGERAKEDDGDGDDDDVEEEVEEKEGTSSTAFQAEE KRRDAGDPGSDVSEATVDFTLVNGSRVGARSKRIHRLSI NQALMVLTDMVVSSQRQRRLRNDNRMRQQLLSIG |
| *Melibe* | Length: 190<br>Signal pep: N/A<br>Cleavage sites: 5<br>Locations: 19, 49, 54, 90, 150 | MNTDDTNMDAVAGSVGSDRSSDDSRGEAFPADSYPSPN YSPYDSNNIYRSLVYRQAQNDDNDKTNSSPPPLQEIGGQ SAYENNVNNVKERSSNADGEFGQGRRRLRLPGPGTAMA ETTGENDGISDIAPAASSRVASTAATAATAPRQKRWQRLS VNQALMVLTDMVMHEQRQERLRKAEKMRAHLLSIG |
| *Flabellinopsis* | Length: 372<br>Signal pep: 1-29<br>Cleavage sites: 9<br>Locations: 98, 151, 177, 232, 241, 259, 280, 316, 332 | MKTAMTAHTNSLLGIFVIAMAATISPILATSPSSYSLSNM ASNMAAHKATSLSRLITDIKSVRLIHNCREFVQQFKNAQ PSSSSSSNKLQHLSKVDIRDVVKNSYTETRIPHFLRHNEIE NNANVDSMLDFKFPTTTEFPLYQNDALTGGKSLPFGYHF ARESSHEKNNNKFPVFQRQDEVKIEKNHFQFMEQMCLK YAFDKLLQTQQKERRRVDSSSDNFMHNNHFNPSERSIPE SVASKIGDEGYILDSNNPHRLSRSRPAYNDGFNTRFNSNN NLPRSSSSPSSQSTSFRSQEMSDNDQQSSDDSSPITPSKRG YSSFANWMRQVRAKRRHRISINNALMTLTNSLVEDRQQ RVKQNRNNMMYHMYLQG |
| *Hermissenda* | Length: 327<br>Signal pep: N/A<br>Cleavage sites: 3<br>Locations: 108, 264, 287 | MAAHSYNDGIIDNTHSFGDNSETSDGFRNKLRNSLHNSK SSTASDDIYPGFYVENTNRFPVHSTDGINEIELKPAKDAF EFMEQMCLKYAFDKLLQSKNRERYWASKKALEETNDV GTEYYDRDSRRPFSGMFGQQQQEDDEDEDDQLLSMTSH PINLDDISIENRPFLSTRLSLSPDYLTYDDGLKSMSNLYNN NYNNNNNPFLNPSSFQEQESEDDNDVMNDRDGLSENTA ARMQLHQQTPDFYDDDDDGDETTEMMSPFKRGGEAPS STMLARLIGGVQTRGKRRHRMSINNSMMTLSSMIWEKN NQRKQKARNNIMRYMQEQG |
| *Pleurobranchaea* | Length: 282<br>Signal pep: 1-24<br>Cleavage sites: 8 | MSPSCRLLASCFFLLSLHVPGCPASPLSGQSYPPMTEAFC RSLLRQIAHADSRSRHRADLDDSEQPISTADFYHVARQS HLPGQISQLAQDFRQDLLDSMATTAFPLTQPADHQTEEA EADRFLSEMSRVPTRCLLHVVGLARLSDAFPPGHGARQ |

| | | |
|---|---|---|
| | Locations: 41, 45, 53, 133, 214, 241, 256, 262 | AGGQGGYSVLGGALGTADIPTPKTEEFGLSMPSSLLGTQ<br>GAKMADGDVTQGDLTSSKRAYRGKSTSKSASKPRWMT<br>SPRHAMSKRPHHRMSINNAFVNLKDMLVARSRQRVMR<br>NRANMRYLMYAQG |
| *Armina* | Length: 348<br>Signal pep: 1-30<br>Cleavage sites: 7<br>Locations: 85, 150, 162, 244, 255, 266, 308 | MLEQIPSVLNIVVMATTMAAIATSSPLTSGSPFFTSLAATK<br>DSKPDSTIPYNLDSDLNPVSLAKECLDYFEQLGPPWSGT<br>DVNNRENPQKESHNIHNPSFNEQPVDSLEEIENHNANSL<br>YPNINSLHTIPNFNSPIGNFPADNEFIKTKRYSQNSLEKMF<br>KRNEGFPNDDKGDLKETTIHILNQICLEFAFSGQPSIRYAN<br>EPSTGIDDKANKFRLAVDVASGPTSTSVHQYMNIAPLTSD<br>PSRISPLLTSHPKRYEYYIYNTMKRSPTSIHNRWNSNRPM<br>GISHPVTSSNRGSSAWLQLNRRMRSKRHHRLSINNSMM<br>TLSNMMIQNNRQRMLKNRAKMMYHMYLQG |

**Supplemental Table 3:** Summary data of nudipleuran gene annotations used to make the gene models in Figure 1

| |
|---|
| ATGAAAACGCCTTTAACCAATTTCATTTTTGACCTCGCCATTATGACAACAACAATGGTT<br>GCTATTATCTCGCCCGTTCACGCAACCTTACGTTCACGAGATCTATTTGCAGACAAATCA<br>TCTCCCCTGCCGAATCTCATTTCAACCATCAAATCGCTACGATTGATGAACACTTGCAGG<br>CAGCATGTCGGACAATGGAAATACAATCCAAATCCGTCTCTGTATTTCCCCTCAAGGACA<br>TTTCAAAACGTCAACCAAAATAGCATCAGAGAAATGGCTAAAACTACAAGTTATAACGAC<br>GCCAATTTCGCAAGCAAACTTATAAACGTTATAAAAAACTCAAAGTCATCAACATCCAAC<br>TTGCCTTTATTTCAAGACGCTTCGTTTTTGGATAACATCTCAGTTGATCTTCGTAAGATG<br>CTAAACGAAAAATTGAACAAAAAGTTCTCTTTATATTCTCACAGCACCAAAACGGATCTT<br>AATGAACAGTCCAATAAAAACTCATTTGAATTCCTGGAGCAAATGTGTCTGAAATACGCT<br>TTTGACAAACTTTTACAAAATCAAAACAGAGAGCAGTATTTGGCTGGCAAAAAAGCTTTA<br>GAGGACAGGAATCATATTGTAAGAAATCTACAGGGTCTACAACAGTTTCCCGAGATGCTT<br>TCACCAGAAACCAGTGGAAACGTCGGCCTTCTATCACTACTTTCTCCTGATGTCAATATC<br>AGAGAATTTCGCCATTCATTTCATATGTCAACTCCTCCTAATCATCCTGTATACAACGAC<br>GGGCGTAGCTTCTCAAATACCTACAATAGTAATCAGAAAATGGAACGATCGCCATTTGAT<br>GCATCCGAAGGAAATATAGTGCCAAGCAGTCTTGCCCAAATCGATCCAGAAGTGCAACAA<br>CAAGATCGTTCACTGGCTCCCAAACGGTCTAAACTGATGGCACCATCAAAACGTGGGTTT<br>CCTTCCTTGTTCACCTGGCTGGGTAATTTCCGGAACAAACGACGACATCGCCTTTCCGTC<br>AACAATGCGATGATGACCTTGTCCAGCATGGTTATGGAAGATGCAAGACAACGAGCCCAG<br>AAGAATCGGAATAACATAAAATATCACATGTACCTGCAGGGGTGA |

**Supplemental Table 4:** *Berghia stephanieae* ELH preprohormone full nucleotide sequence as used to make HCR probes.
